# Computational approaches towards reducing contamination in single-cell RNA-seq data

**DOI:** 10.1101/2020.07.15.205062

**Authors:** Siamak Yousefi, Hao Chen, Jesse F. Ingels, Melinda S. McCarty, Arthur G. Centeno, Sumana Chintalapudi, Megan K. Mulligan, Pete A. Williams, Simon J. John, Bryan W. Jones, Monica M. Jablonski, T J. Hollingsworth, Eldon E. Geisert, Lu Lu, Robert W. Williams

## Abstract

Single cell RNA sequencing has enabled quantification of single cells and identification of different cell types and subtypes as well as cell functions in different tissues. Single cell RNA sequence analyses assume acquired RNAs correspond to cells, however, RNAs from contamination within the input data are also captured by these assays. The sequencing of background contamination as well as unwanted cells making their way to the final assay Potentially confound the correct biological interpretation of single cell transcriptomic data. Here we demonstrate two approaches to deal with background contamination as well as profiling of unwanted cells in the assays. We use three real-life datasets of whole-cell capture and nucleotide single-cell captures generated by Fluidigm and 10x technologies and show that these methods reduce the effect of contamination, strengthen clustering of cells and improves biological interpretation.

## INTRODUCTION

Single cell technologies have made numerous advancements in understanding cell types and behavior. ^1-3^ Efforts are being made towards generating human cell atlases.^4^ BRAIN Initiative is another example of such efforts with the same goal; identifying different brain cells. However, all these efforts are highly dependent on the accuracy, reproducibility, and repeatability of experimental and computational workflows of single cell RNA sequencing (scRNA-seq) transcriptome experiments.

A fundamental assumption underlying scRNA-seq data analysis workflow is that data contain only mRNA from single cells and single cells belong to only the tissue of interest. However, both assumptions may be violated in single cell experiments. For example, doublets and triplets, empty droplets, and other cells than the targeted cell may exist in the data. Removing contamination both experimentally and computationally are active research areas.

Here we show that contamination exist in three real-life dataset that we have generated from retinal ganglion cells (RGCs). RGCs essentially carry visual signals from the retina to the brain.
^5 6^ While all RGCs share a common characteristic of lengthy axons, their physiological roles in responding to visual stimuli may differ. Different RGC types transmit different information to the brain, such as changes in the intensity of the light (ON/OFF cells) or moving objects in a specific direction.^7, 8^ Traditionally, RGCs have been classified to different types based on their morphological and physiological characteristics. ^9 10^ Identification of RGC types is critical to understanding the mechanisms underlying retinal diseases such as glaucoma. Moreover, it promotes the reproducibility of conducting experiments at different laboratories provided all working on a specific RGC type. However, to date, the identity and number of RGC types is unclear. ^11-13^

In mice, advancement of the genetic techniques have identified over 30 different subtypes of RGCs based on molecular markers. ^1-3^ A hypothesis (used in other neural systems) is that different RGC types express specific sets of transcription factors and as a result drive genes that are important for different type-specific features. ^14^ To identify different RGC subtypes using molecular markers, we generated two scRNA-seq and one snRNA-seq datasets and realized that all data includes different types of contamination. In this work we focus on non-RGC contamination and how to mitigate these sources of contaminations using domain knowledge and computational approach.

## METHODS

Briefly, our experimental protocol consists of the following steps (Figure 1): (1) removing the retinas from 130- to 150-day-old mice, (2) trimming the optic nerve head and pooling retinas, (3) dissociating gently and enriching using THY1 antibody-coated micro particles (beads), (4) isolating and generating single cell RNA libraries of full length polyA-positive mRNAs, (5) sequencing the libraries. The computational workflow included the following steps (Figure 2): (1) loading data of single RGCs into R program, (2) quality control and excluding duplicate genes, (3) filtering cells and genes, (4) normalizing data and log-transforming, (5) selecting highly variable genes, or alternatively, using domain knowledge to select genes that are known or candidate markers for RGCs but not known markers for other retinal cell types (e.g., Amacrine, Bipolar, Microglia, Rods, Cones and Astrocyte cells), (6) linearly reducing dimensions using principal component analysis (PCA), (7) non-linearly reducing dimensions using t-distributed stochastic neighbor embedding (tSNE), (8) applying unsupervised clustering to identify clusters (this step can be performed after step 6 too), (9) performing hierarchical clustering to identify retinal cell families, (10) apply differential gene expression to find biological annotations for cluster of cell types.

**Figure 1.**
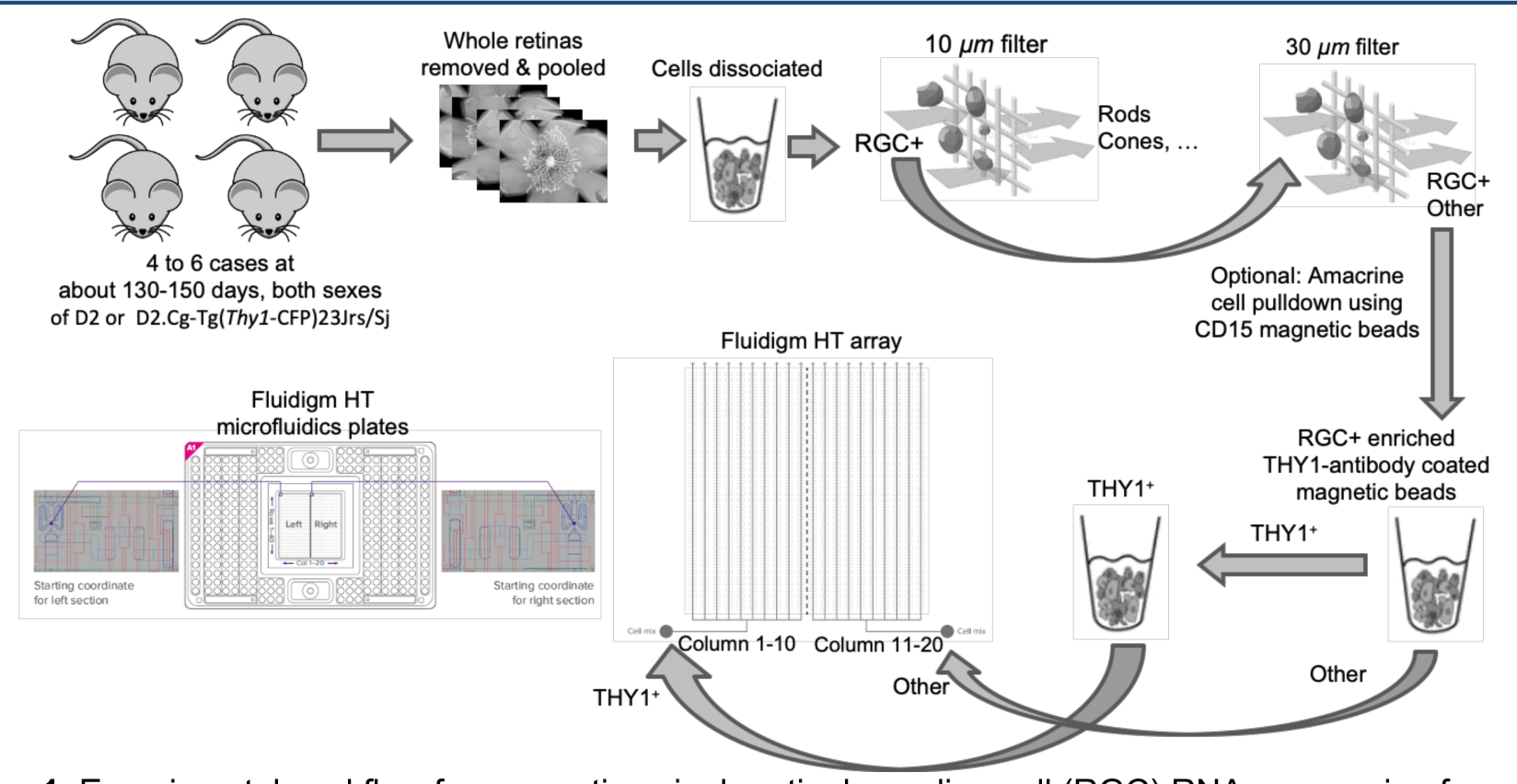
Experimental workflow for generating single retinal ganglion cell (RGC) RNA sequencing from retinas of glaucoma mice using Fluidigm technology.

**Figure 2.**
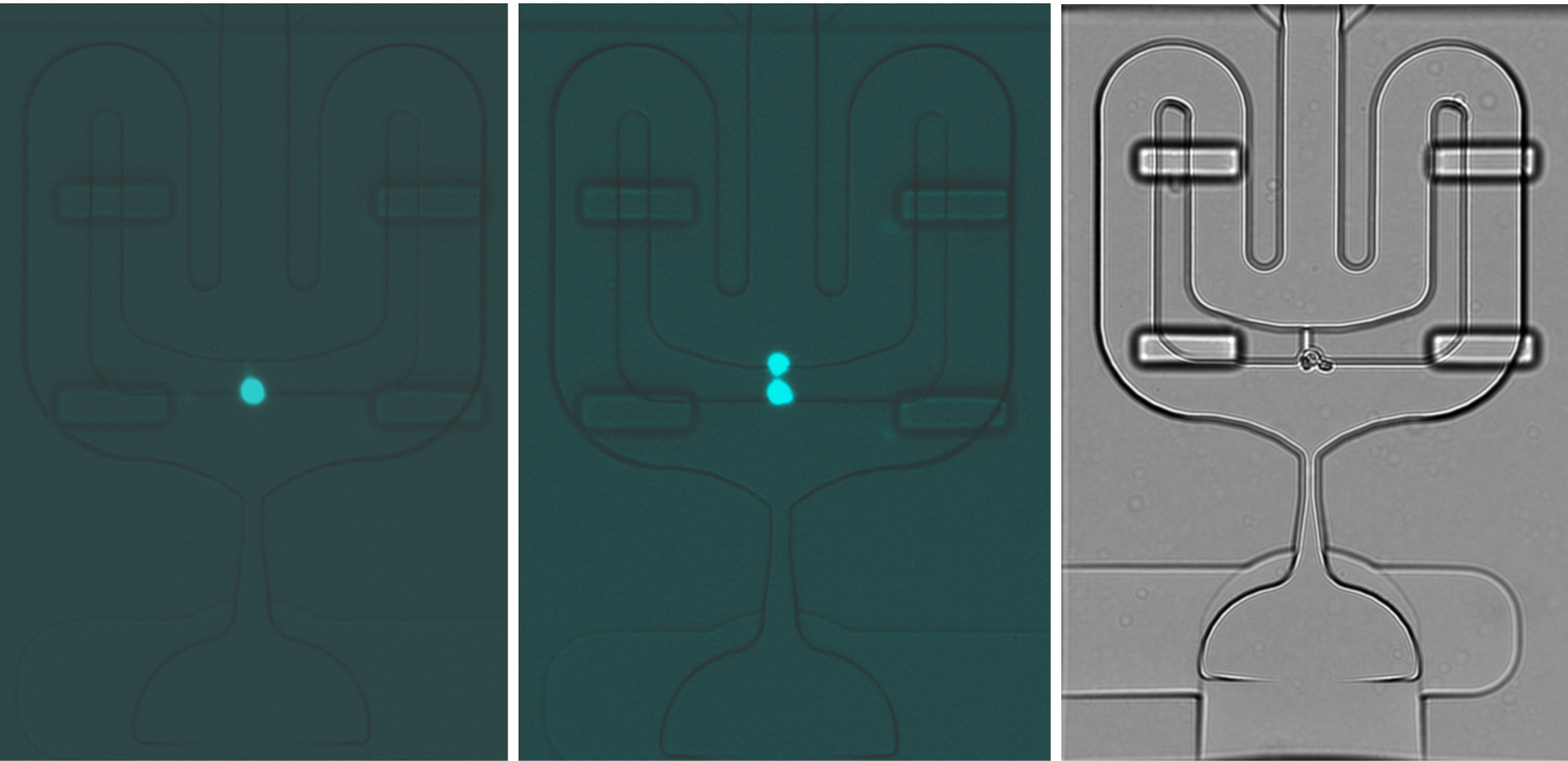
Images from single cells in Fluidigm arrays. **Left** is an example of a perfect cell, middle represents two cells and right panel shows three cells that entered the Fluidigm array.

### Samples generated by Fluidigm technology

Four to six DBA/2j or D2.Cg-Tg (thyl-CFP)23 Jrs/Sj mice in an age range of 130-150 days old were anesthetized and the retina were removed in one piece (Fig. 1). The optic nerve was trimmed away, and the harvested retina are split between two tubes. The two tubes of pooled retina were given a short spin to collect the tissue to the bottom of the tubes. The tissue was then mechanically dissociated by pipetting tubes up and down. Each cell strainer was pre-wetted and given a short spin to collect the contents and the cell suspension all less than 70 µm. The cell suspension is added to a 10 µm Plurifilter allowing all cells less than 10uM to filter through with those greater than 10 µm to remain on the filter. Another 30 µm Plurifilter was added, to wash the cells greater than 30 µm. Then RGCs were enriched using THY1 antibody-coated beads. Fluidigm HT microfluidics plates were used to isolate and generate single cell RNA libraries of full length polyA-positive mRNAs using SMART-Seq v4.

While we expect only a single cell in each Fluidigm array, in practice, doublets, triplets, or debris enter the array. Figure 2 shows three sample images from cells in our Fluidigm array captured by Nikon Eclipse Ti2 microscope.

### Whole single samples generated by 10x technology

Briefly, 11 DBA/2J mice, age range 130-159 days old were selected for 10x whole cell Isolation. Intact retina was removed, and dissociation of retina was accomplished using the Neural Dissociation Kit (P) (Miltenyi Biotec). The two tubes of pooled retina were given a short spin to collect the tissue to the bottom of the tube and the media was completely removed. The 10uM Pluri-filter was inverted onto a pre-wetted 30uM Pluri-filter, inserted in a new 50 ml tube and 5mls of cold HBSS W/O was added (1ml at a time) to wash the cells greater than 10uM onto the 30uM Pluri-filter below to allow the cells less than 30uM to flow thru into the 50 ml conical tube while the cells greater than 30uM remain on the filter. Cells with beads attached remain in the column and the flow-thru should contain non-THY1/CD90.s+ cells and debris were discarded. CD90.2 labeled cells flushed thru the column with a plunger after the addition of 1.5mls MACS Buffer. This tube containing THY1/CD90.2+ whole cells was filtered thru a 30uM filter then washed one time with HBSS and the pellet re-suspended. An aliquot of the CD90.2+ whole cells were counted on the Countess II (Life Technologies) using Trypan Blue exclusion dye and the cells diluted to a target concentration. Figure 2 shows the overall 10x workflow for generating single RGCs. A total of 5,820 whole RGCs with a mean of 142,609 reads per cell and a median of 3,523 genes per cell were generated. Over 99.9% of the UMIs were reported as valid by cell ranger.

### Nuclei single samples generated by 10x technology

Characterization of phenotypic diversity is an active yet challenging field of scRNA-seq. Patterns of gene expression are typically used to explore single cell heterogeneity. However, patterns of gene expression are subject to a different source of variation and noise such as batch effect and cell cycle. One approach to cope with these sources of variations is to use single nucleotide rather than single cells data.

To that end, we generated single nucleotide cells to explore RGC subtypes. All steps above were followed to isolate CD90.2+ whole cells and once whole cells were isolated, a Nuclei wash, and re-suspension buffer were prepared to isolate nuclei. Suspension was pipetted up and down 5 times with a wide bore 1000ul pipette tip. Cells were centrifuged and the supernatant was removed without disturbing the nuclear pellet. The nuclei were re-suspended in Nuclear Wash and Re-suspension Buffer, centrifuged again. This nuclear wash step was repeated, and nuclei were re-suspended in Nuclear Re-Suspension buffer. The isolated nuclei were passed thru a FlowmiTM Tip Strainer and counted on the Countess II. Nuclei number was adjusted to 1000 nuclei/ul and held until the 10x Genomics run. Figure 3 shows the overall workflow for capturing single nuclei RGC nuclei. A total of 10,000 nuclei with 222,603 Mean Reads per Cell and a median of 1,613 genes were generated. Over 99.9% of the UMIs were reported as valid by cell ranger.

**Figure 3.**
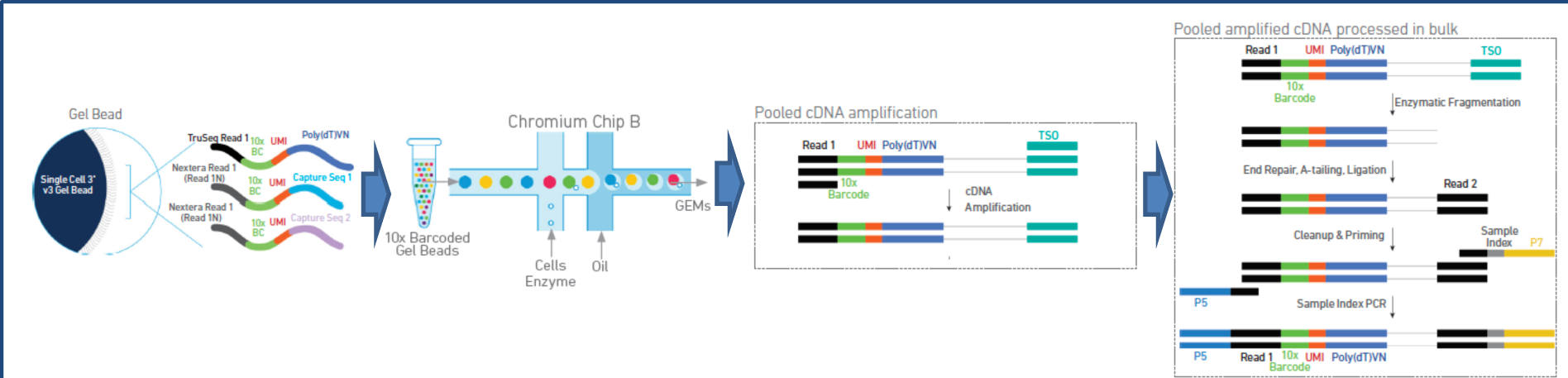
Experimental workflow for generating single retinal ganglion cell (RGC) RNA sequencing from retinas of glaucoma mice using 10x chromium technology.

### Data preprocessing and quality control

In Fluidigm, we sequenced 9,600 cells (the libraries were collected in 12 different plates (batches), each plate included 800 cells) using HiSeq 3000, 150 nucleotide pair-end reads. We de-multiplexed barcode rows using R1 reads and timed and aligned R2 reads to reference genome to generate gene expressions. Data was loaded into R software. All plates were merged based on unique gene names. We excluded two plates due to technical issues. We generated a big transcriptome data of size [43,320 transcript models * 9,600 cells]. We first performed preprocessing and applied quality control. Two plates had technical issues and were removed from the analysis. To exclude duplicated genes from data, we selected genes with greater number of non-zero expression across remaining cells. In another word, we selected a gene that was expressed more frequently across cells regardless of the expression level. This step generated 25,394 unique genes. We then selected cells that expressed at least 900 genes and selected genes that had been expressed in at least five cells. A total of 16,622 genes and 6,222 cells were selected for the downstream analysis. This step reduces the change of including fragments in the analysis.

### Normalization and linear transformation

We then normalized the data; for each cell across all genes, we normalized the Fragments Per Kilobase Million (FPKM) values to the total expression using functions in the Seural package.^15, 16^ We transformed the data to the log2 scale. We first applied principal component analysis (PCA) in order to ensure robust identification of the primary structures in the expected single cell data. PCA uses a linear and orthogonal transformation to convert the observations of highly correlated transcriptomes into a set of new eigen-genes (principal components) which are linearly uncorrelated to each other. In another word, each new eigen-gene is a weighted combination of all initial transcriptomes while the eigen-genes do not carry correlation anymore.

PCA transformation allowed us to linearly reduce the number of dimensions of the original dataset. While the number of identified eigen-genes is equal to the number of input genes, only a small fraction of eigen-gens can explain a significant portion of the variance in the data. We used JackStraw method, which is a randomization approach by creating a null distribution of the eigen-genes identified by applying PCA to 1,000 new realizations of the input data in which we randomly scrambled 2% of the genes. We then selected eigen-genes from the original dataset in which their scores were significantly different from the respective scores derived from the null distributions (p<0.01, Bonferroni corrected). ^17, 18^ We then subjectively verified PCA-based markers for distinct retinal cell subtypes from the absolute value of the eigen-gene scores. We note that these scores leverage information from genes in the PCA, and therefore are more relevant to retinal cell subtypes as well as more robust to technical noise than the original data of all gene expressions. We used the selected eigen-genes for the downstream analysis.

### Excluding PCs (eigen-genes) with strong correlation with known markers of unwanted cells to reduce the impact of contamination

This is an intermediate solution to decrease the effect of unwanted contamination computationally by excluding PCs with significant scores of markers from other unwanted cell types; non-RGC markers in our case. For instance, if major scores of a PC is composed on marker genes of photoreceptors in retina, we will exclude that PC from the downstream analysis to reduce the effect of those unwanted cells as one source of contamination in the dataset.

### High-density based clustering to reduce the impact of contamination

After excluding potential eigen-genes with strong scores, another approach to reduce the contamination is identification of clusters of unwanted cells using manifold learning. We used t-distributed stochastic neighbor embedding (tSNE)^19^ to group cells with similar eigen-genes together. Eigen-genes were used as the input for tSNE. This process mapped cells with similar local gene expressions, and therefore similar eigen-genes, localized in the tSNE space while nonlinearly reducing the dimension of the eigen-genes to two-dimensional embedding of single cells, and hence distinct cell types formed two-dimensional clusters. Moreover, tSNE provided a well-suited visualization of high-dimensional transcriptome data with outcome in 2-dimensional tSNE space.

To identify cell types in the tSNE space (putative RGC subtypes here), we employed a density-based clustering to identify cells with similar gene expression patterns, and hence similar eigen-genes and tSNE scores, and to group cells into non-overlapping clusters objectively. ^20^ In fact, density-based clustering groups those cells in the tSNE space that that are closely packed together and have many neighbors around them while cells that lie alone (in low-density areas) and are too far away will be marked as outlies and non-members of the clusters. We initially set the reachability distance parameter (eps) to 1.6 to over-partition the cells in the tSNE space provided us 35 clusters. This process allowed us to identify and subsequently exclude clusters retaining fewer than 30 cells (∼0.5% of the initial 6,222 cells). Clusters retaining small number of cells were excluded from the analysis. These small clusters typically located along the interfaces of larger clusters. We then repeated the procedure with a larger eps value to identify only well-separated dense clusters of cells. This pruning approach enabled us to exclude outlier cells while avoiding over-partitioning of the cells. We next investigated the partitions objectively to ensure that our identified clusters represented distinct groups of cells, as opposed to merely over-partitioning groups.

We will then perform post-hoc analyses to interpret and assign biological attributes to the identified clusters quantitatively using known markers for unwanted cells. For instance, from literature, we identify known markers for retinal cell types such as Microglia, Rods, Cones, and Astrocytes and try to identify cluster of cells belonging to these groups. We will then exclude those clusters from downstream analysis.

The final step after exclusion of potential clusters of unwanted cells, we identified the differentially expressed genes between every pair of clusters by simultaneously testing the difference in frequency and expression level of genes using likelihood-ratio test^21^ and iteratively merged highly related pairs with the lowest number of differentially expressed genes. None of the clusters met the criterion of less than two differentially expressed genes for merging.

We used R software to develop the above-mentioned workflow. We used tools that were used in previous landmark articles ^22-24^ along with some of the functions and tools provided in the R library of Seurat package. ^16^

## Results

### Fluidigm whole single cell capture

We included 9,600 putative RGC cells generated from 12 sequencing runs of Fluidigm technology and generated a single matrix of size (9,600* 43,320). After exclusion of runs with major technical issue in the workflow process, 7,195 cells entered the analysis. The data was normalized by dividing to the total FPKM per cell. All analysis was conducted in the log2 space. In average 2,934 genes were expressed per cell (Fig. 4). To enhance the power of unsupervised clustering for discovering RGC subtypes, we filtered cells expressing fewer than 900 genes and genes expressed in fewer than five cells. A total of 6,222 cells expressing 16,622 genes were included for the downstream analysis. Figure 4 shows the histogram of the number of genes expressed in cells. In average, 2,934 genes were expressed in cells.

**Figure 4.**
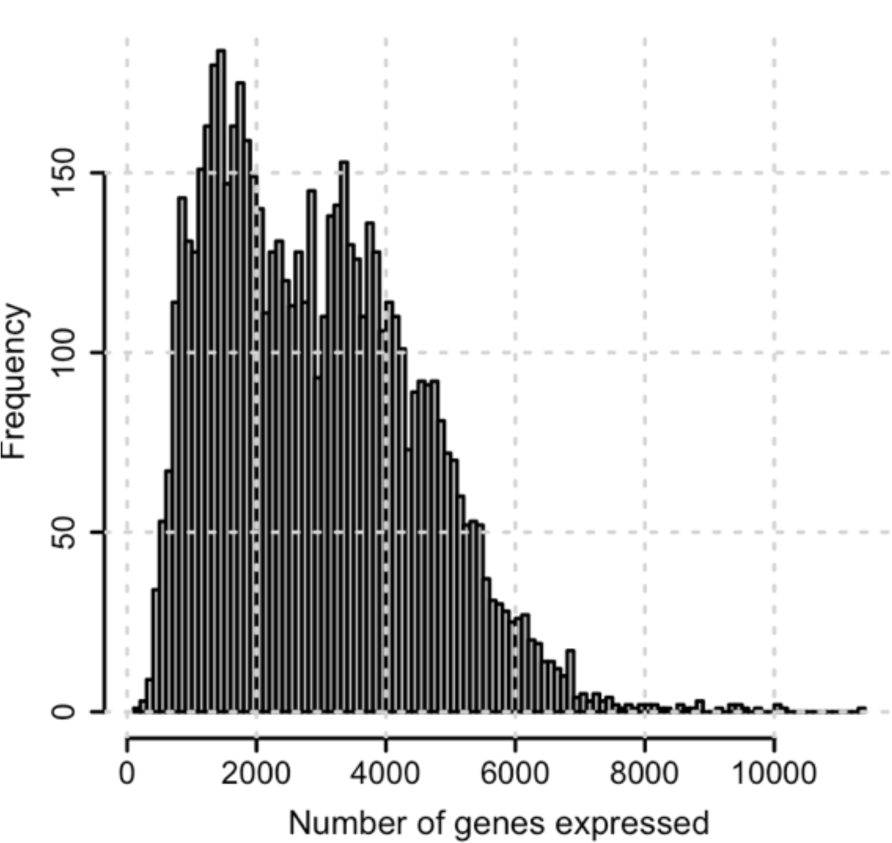
Number of genes expressed in putative retinal ganglion cells (RGCs) whole cell capture by Fluidigm technology.

We then scaled and centered the data along each gene then selected highly variable genes by computing average expression and dispersion of each gene. We identified 2,048 highly variable genes based on average expression and dispersion (Fig. 5). Reducing the number of genes reduces the computational complexity and improves the ability to identify selective genes of different RGC types.

**Figure 5.**
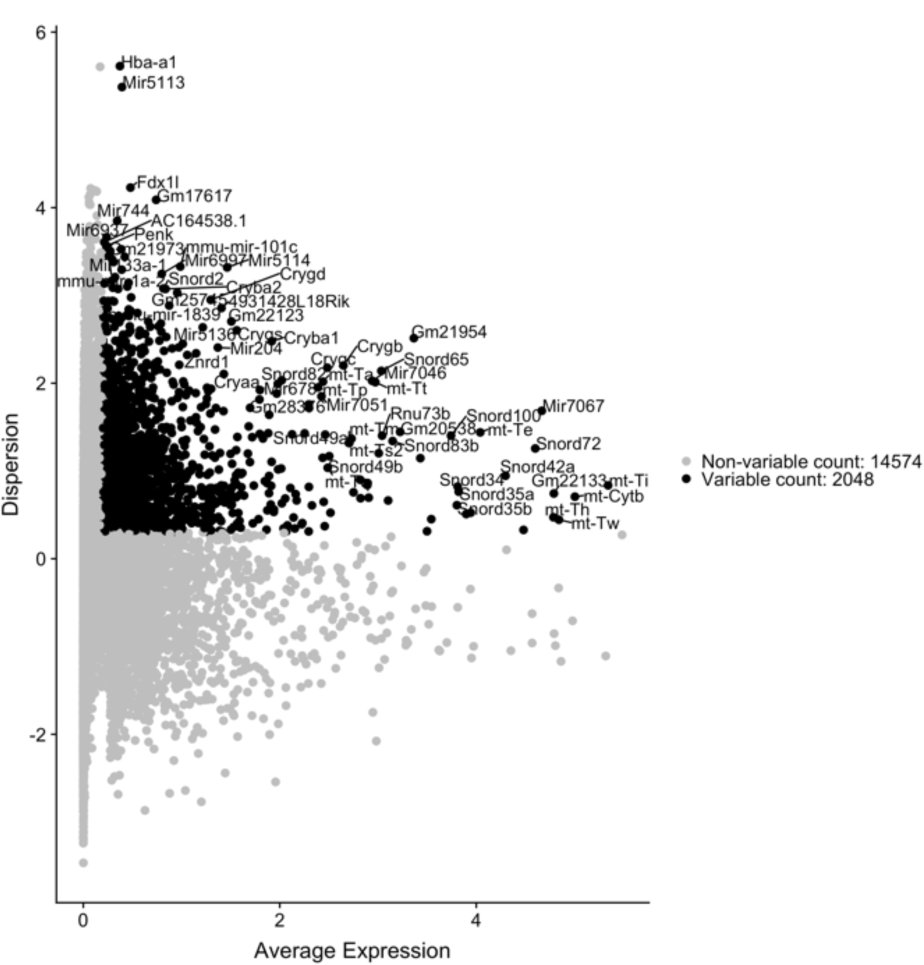
Average expression versus dispersion of genes in our dataset. Genes that having an average expression and dispersion of greater than a threshold are selected (represented in black). Names of some of the highly variable genes are provided.

We then applied PCA on the subset of 2,048 highly variable genes in order to capturing the primary structures and patterns in the transcriptome data. This process generates 2,048 principal components, however, only a small number of these components capture the variance exist in the data. We used the elbow plot followed by JackStraw^7, 18^ method and selected 28 significant PCs for the downstream analyses. Figure 6 demonstrates the data in the principal component spaces. These 28 principal components are more robust to technical noise than individual gene expression values. We computed the scores of genes in these 28 PCs.

**Figure 6.**
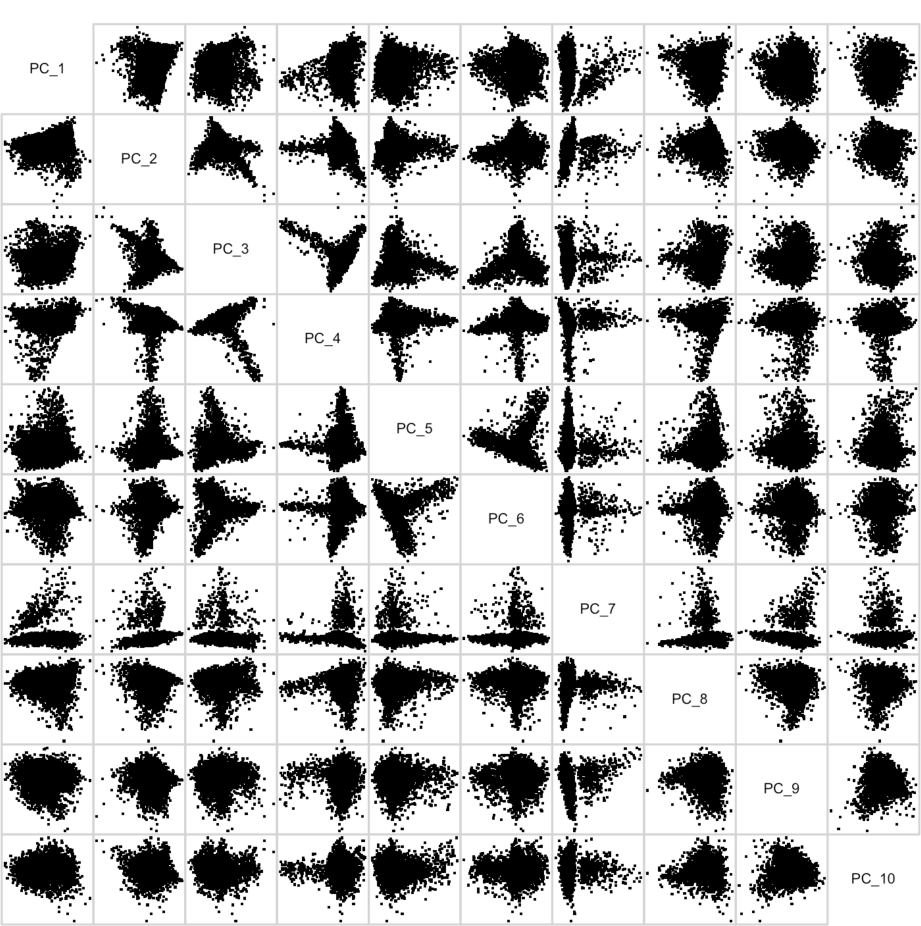
Principal components. Each panel represents data in two of the principal components.

To identify PCs representing non-RGC markers, we found 68 genes (e.g., Gpr37, Gldc, and Epas1) from literature that are known markers for major non-RGC retinal cell types including Muller, Fibroblast, Astrocyte, Amacrine, Endothelium, Horizontal, Photoreceptor, and Microglia cells. We then computed the pairwise correlation between any of these 68 markers and any of 40 PCs. We then excluded those PCs that showed a significant correlation (Pearson) with known non-RGC markers. Second PC had the highest correlation of 0.35 with a few non-RGC markers and was excluded from the downstream analysis. This step requires domain expertise and essentially integrates biological knowledge into the computational workflow but could be skipped if appropriate domain knowledge is lacking.

We then performed graph-based clustering that starts with a K-nearest neighbor (KNN) graph, with edges drawn between cells with similar gene expression patterns, and then partitioned this graph into highly interconnected quasi-cliques or communities, as outlined in previous publications.^25 26^ We then applied modularity optimization techniques proposed in Louvain algorithm or SLM ^27^, to iteratively group cells together, with the goal of optimizing the standard modularity function. We visualized the clusters using tSNE subsequently. ^19^ We set the effective number of neighbors in tSNE represented by “perplexity” parameter to 30. In fact, cells with similar gene expression patterns will fall closely, and hence distinct cell types should form clusters in the 2-dimensional tSNE space. Figure 7 shows the tSNE plots of the cells we identified. Left panel show mapping of genes onto tSNE space. In the right panel, cells are color-coded by their plate. We propagated the cluster labels identified by SLM unsupervised learning and color coded and numbered the clusters, as shown in Figure 8.

**Figure 7.**
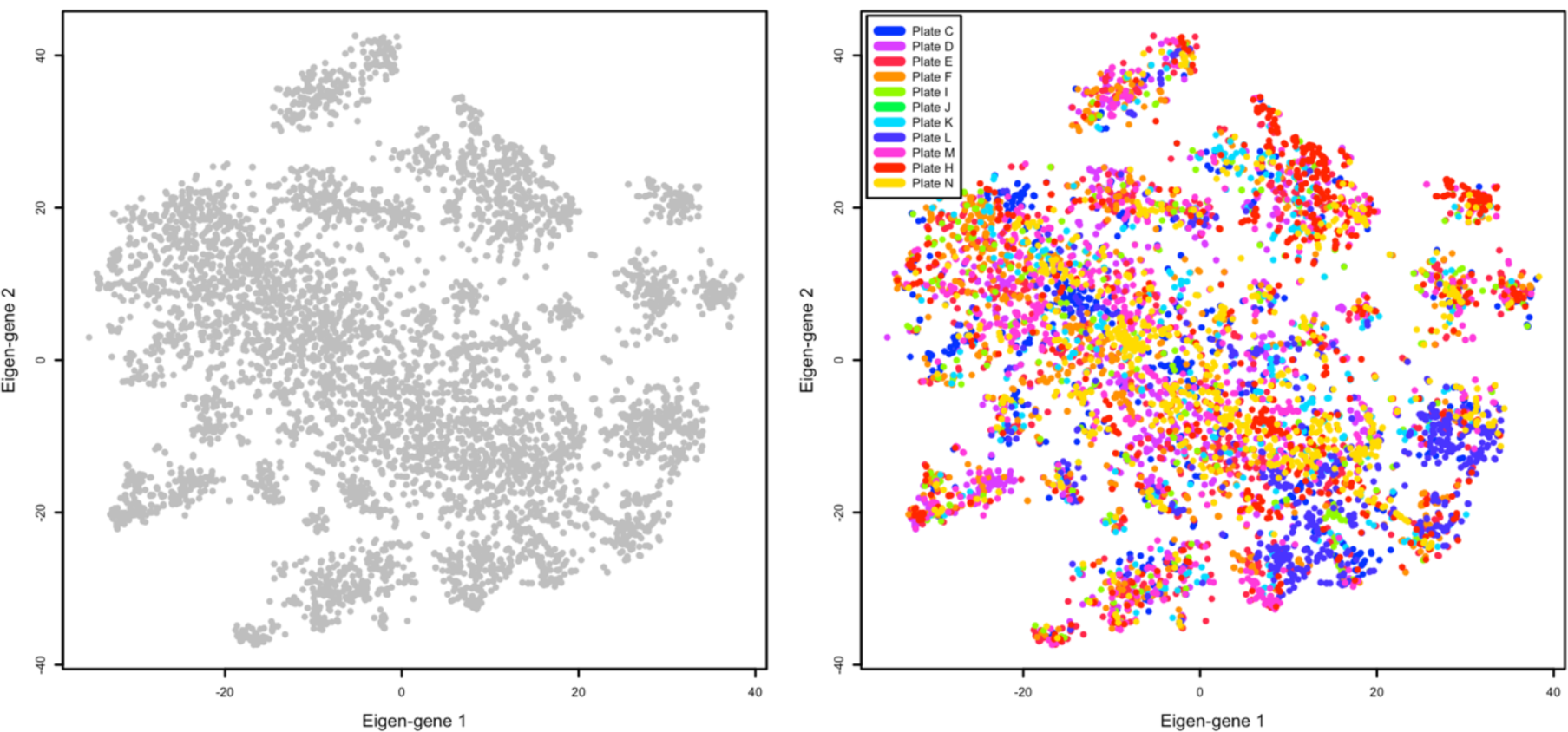
Cell mapped on 2-D tSNE space. Cells with similar transcriptome patterns have been grouped together. Left panel shows all cells and right panel shows cells color coded by the corresponding plate (batch) in the experimental workflow.

**Figure 8.**
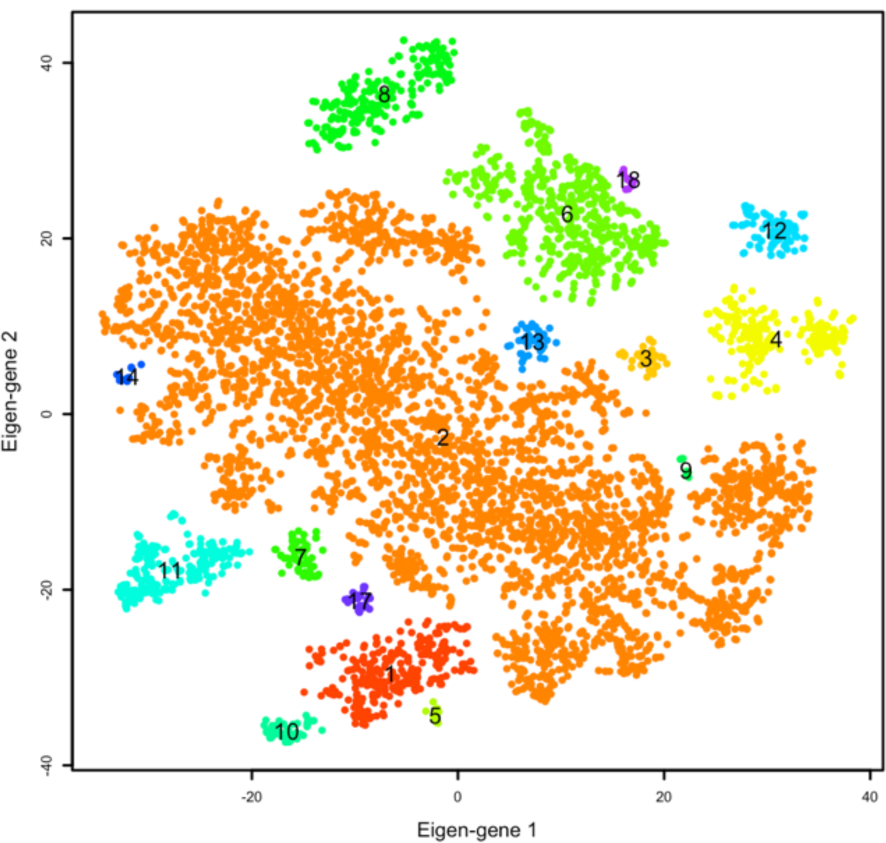
Final clustering to identify cluster of contaminating non-RGC cells.

To identify cluster of non-RGC cells in the tSNE space, we identified marker genes differentially expressed in each cluster by comparing genes expressed in each cluster with genes expressed in the remaining clusters.

More specifically, we required a gene in each cluster to be detected at a minimum percentage and a minimum expression level compared to genes in the rest of clusters. This was investigated using likelihood-ratio test for single cell gene expression.^21, 28^ We explored other techniques including negative binomial generalized linear model, and negative binomial distribution implemented in the DESeq2 algorithm^29^ to assure repeatability. We then performed supplementary *post hoc* qualitative analysis to identify biological attribute of clusters, particularly those with significant expression of non-RGC markers. Specifically, we identified clusters of cells that co-express known non-RGC markers indicated before. This is a classical approach and has been used in the latest studies aiming at identifying different cell types.

We identified that 14 out of 33 genes differentially expressed in cluster number 14 were known photoreceptor markers. Therefore, we suspect cells in this clusters were mostly photoreceptor cells or cells carrying fractions of photoreceptors RNA thus excluded all cells in this cluster from the analysis. This process, coupled with domain knowledge, can be performed iteratively to exclude other potential non-RGC cells from the analysis.

We also performed histological validation. Briefly, eyes from DBA2/J mice were fixed and subsequently dehydrated, cleared, and infiltrated. Then 8 mm sections were cut, and sections were deparaffinized, heated, washed, and subsequently blocked in. Primary antibodies were applied, and slides were washed free of primary antibody to stain the nuclei and incubated subsequently. Slides were then washed and mounted then were imaged using a Zeiss 710 LSM at 200X magnification. Figure 9 shows the plots of major genes expressed in ganglion cell layer (GCL) of the retina verified by histological validation.

**Figure 9.**
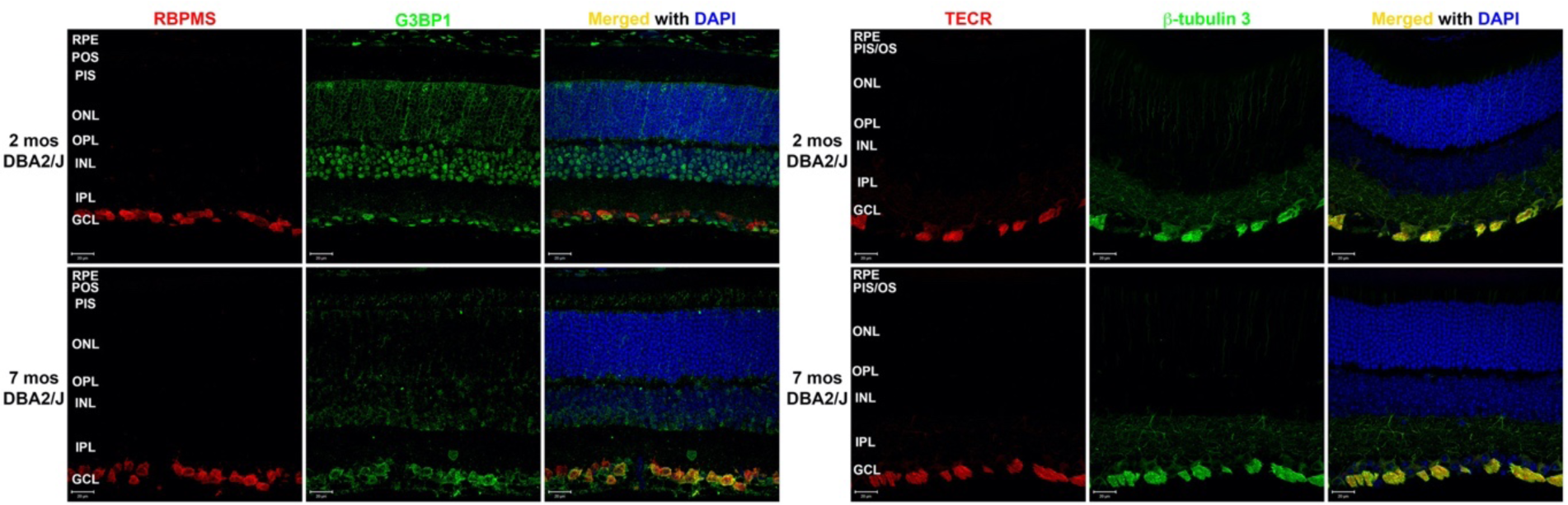
Histological validation of RGC markers. RBPMS and TECR genes were expressed selectively in the ganglion cell complex layer of retina reflecting agreement with our computational analysis.

### 10x chromium whole single cell capture

We purified RGCs from the left and right eyes of 11 DBA/2J mice of age range 130-159 days old by C1 system of Fluidigm technology utilizing microfluidic and CD90.2 magnetic microbeads for RGC surface markers. Microfluidic technology consumes smaller volumes of reagents compared to successors and can automate downstream RNA processing reactions for sequencing. After purification, we immediately processed RGCs using 10x Genomics Chromium platform (Figure 3). ^30^ Each cell was sequenced to the depth of ∼142,000 with an average of 3,216 genes expressed per cell. Over 99.9% of the reads had valid UMIs. Figure 10 shows the histogram of the number of genes expressed in cells.

**Figure 10.**
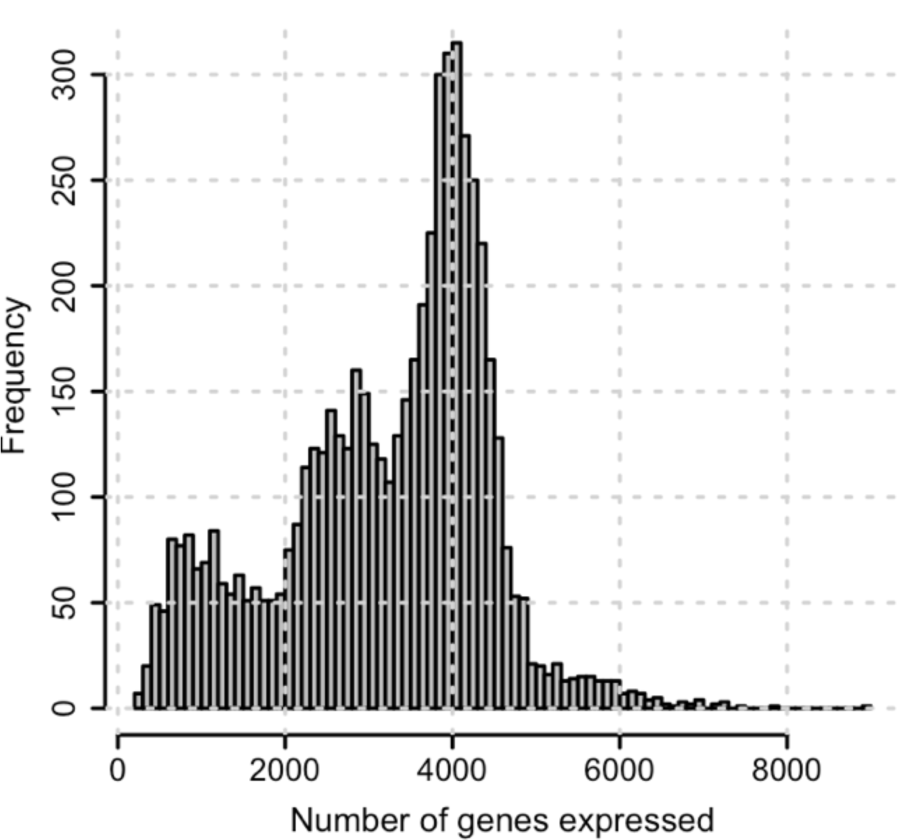
Number of genes expressed in putative retinal ganglion cells (RGCs) using whole cell capture by 10x technology.

To enhance the power of analysis, we filtered cells expressing fewer than 200 genes and genes expressed in fewer than five cells. A total of 5,813 cells expressing 17,997 genes were included for the downstream analysis.

We then scaled and centered the data along each gene then selected highly variable genes by computing average expression and dispersion of each gene. We identified 5,396 highly variable genes based on average expression and dispersion (Fig. 11).

**Figure 11.**
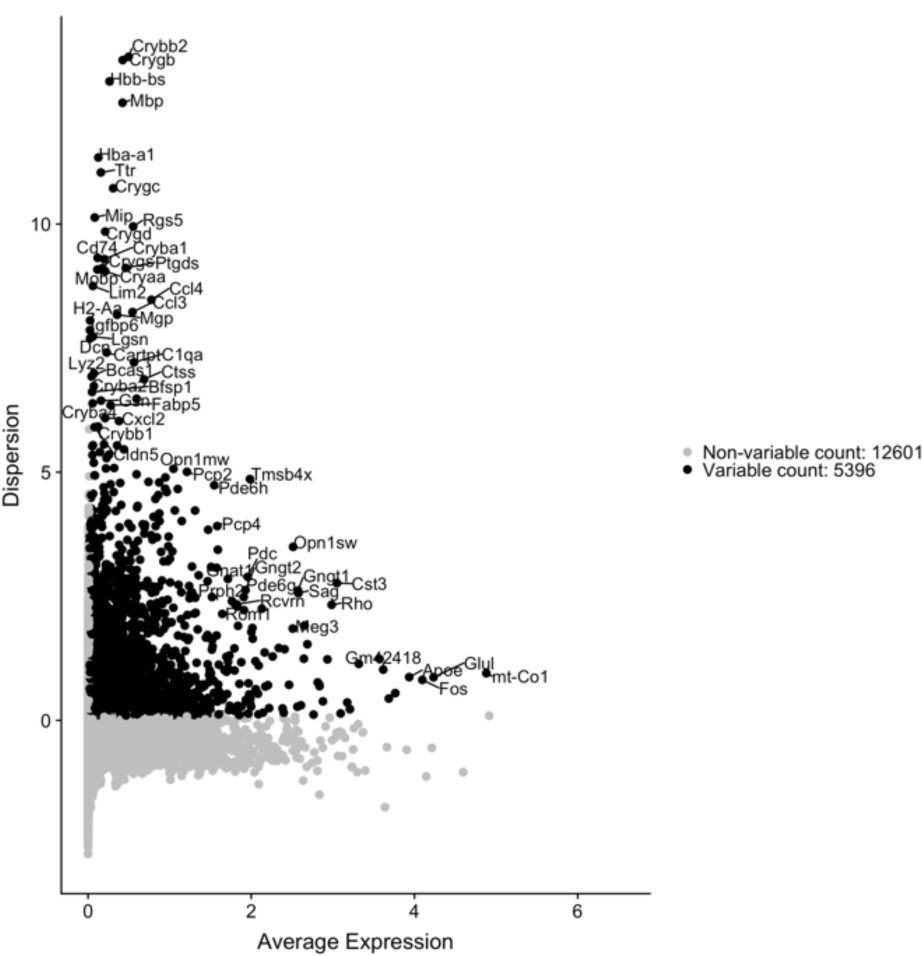
Average expression versus dispersion of genes in the whole cell capture generated by 10x technology. Genes that having an average expression and dispersion of greater than a threshold are selected (highlighted in black).

We then applied PCA on the subset of 5,396 highly variable genes and generates 5,396 principal components. JackStraw^7, 18^ identified 40 significant PCs. We then computed the scores of genes in these 40 PCs. To identify PCs representing non-RGC markers, we found 68 genes (e.g., Gpr37, Gldc, and Epas1) from literature that are known markers for major non-RGC retinal cell types including Muller, Fibroblast, Astrocyte, Amacrine, Endothelium, Horizontal, Photoreceptor, and Microglia cells. We then computed the pairwise correlation between any of these 68 markers and any of 40 PCs. We then excluded those PCs that showed a significant correlation (Pearson) with known non-RGC markers. Second PC had the highest correlation of 0.72 with a few non-RGC markers and was excluded from the downstream analysis.

Graph-based clustering^25,26^ with SLM modularity optimization^27^ identified 32 clusters visualized using tSNE.^19^Figure 12 shows the tSNE plot of the identified clusters color-coded and numbered by labels from SLM unsupervised learning.

**Figure 12.**
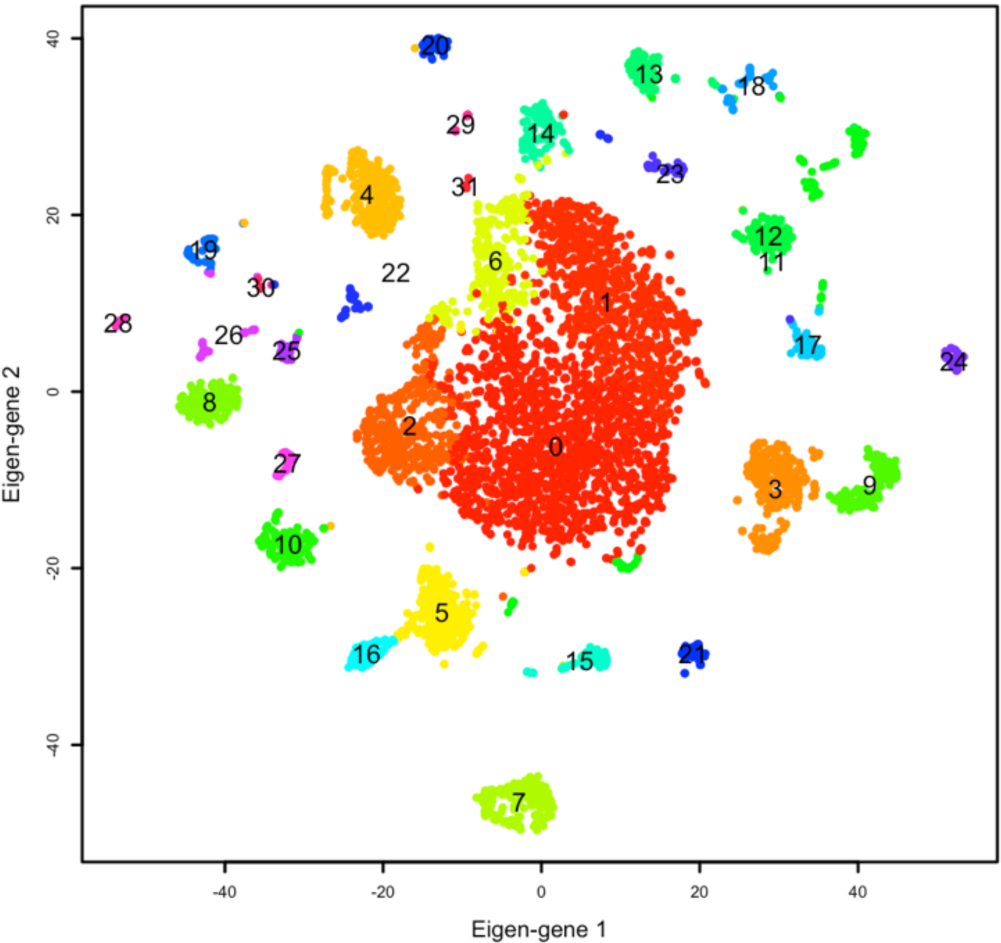
Final clustering identified 32 clusters of cells. Contaminating clusters of non-RGC cells will be identified and excluded.

To identify non-RGC cells in the tSNE space, we first identified marker genes differentially expressed in each cluster by comparing genes expressed in each cluster with genes expressed in the remaining clusters using likelihood-ratio.^21,28^ Post hoc analysis to qualitatively and quantitively identifying biological attribute of clusters, particularly those with significant expression of non-RGC markers, identified a few closers of non-RGC cells.

Figure 13 shows the heatmap of genes that were differentially expressed in 32 clusters in a single snapshot and figure 14 presents sample genes that were highly selective in terms of expression in different clusters.

**Figure 13.**
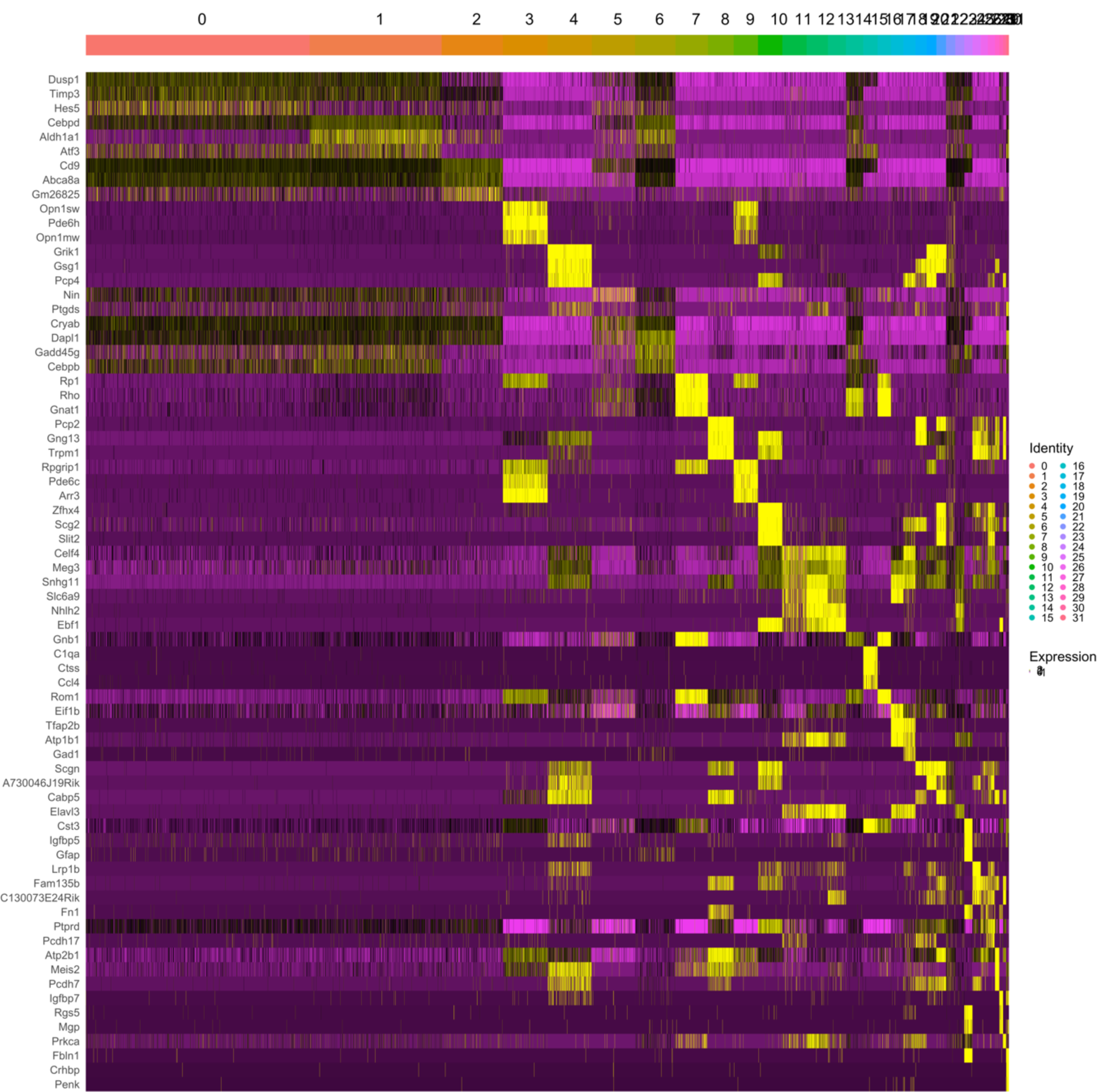
Heatmap plot of genes expressed in different clusters. Cells are sorted based on cluster identity as shown on top of the plot. Genes that were differentially expressed are presented on the y axis. Yellow color shows high expression.

**Figure 14.**
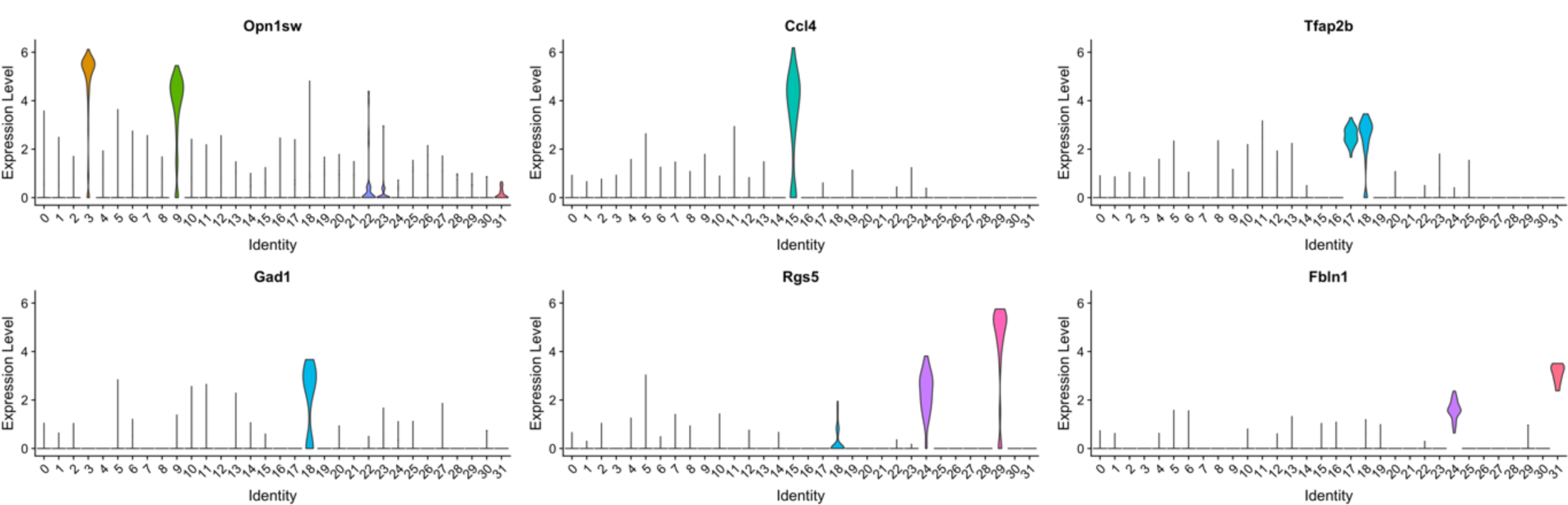
Differentially expressed genes. Six sample differentially expressed marker genes for several clusters computed suing likelihood-ration.

After posthoc analysis, we identified that 27 out of 60 genes differentially expressed in cluster number 7 were known photoreceptor markers. Therefore, cells in this cluster were excluded from the analysis. This process, coupled with domain knowledge, can be performed iteratively to exclude other potential non-RGC cells from the analysis.

### 10x chromium single nuclei capture

We purified RGCs from the left and right eyes of 11 DBA/2J mice of age range 130-159 days old by C1 Fluidigm system utilizing microfluidic and CD90.2 magnetic microbeads for RGC surface markers. Microfluidic technology consumes smaller volumes of reagents compared to successors and can automate downstream RNA processing reactions for sequencing. Once whole cells were isolated, we isolated nuclei by removing the supernatant without disturbing the nuclear pellet. The isolated nuclei were passed thru a strainer and counted on the Countess II software. Nuclei number was adjusted to 1000 nuclei/ul. We then immediately processed RGCs using 10x Genomics Chromium platform (Figure 3).^30^ Each nuclei was sequenced to a depth of ∼220,000 with an average of 1,728 genes per cell.

To enhance the power of analysis, we filtered cells expressing fewer than 1,200 genes and genes expressed in fewer than five cells. A total of 9,442 cells expressing 16,577 genes were included for the downstream analysis. We then scaled and centered the data along each gene then selected highly variable genes by computing average expression and dispersion of each gene. We identified 3,792 highly variable genes based on average expression and dispersion (Fig. 15).

**Figure 15.**
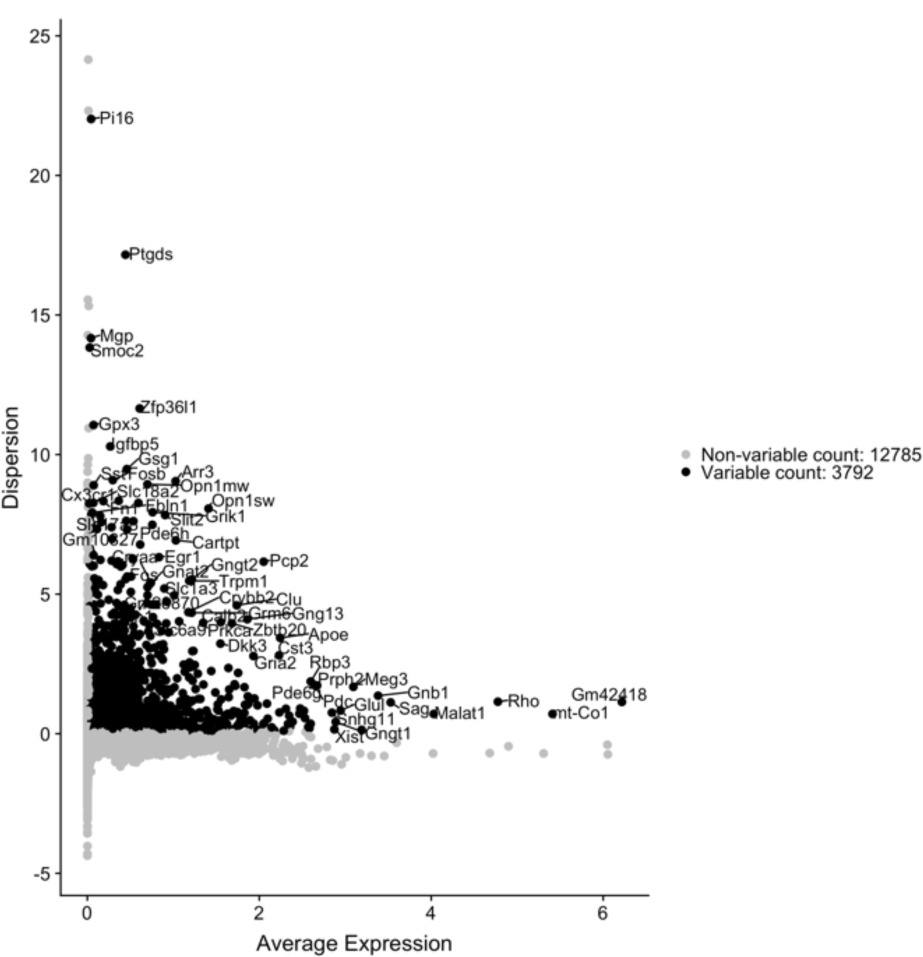
Average expression versus dispersion of genes in the nuclei cell capture generated by 10x Chromium technology. Genes that having an average expression and dispersion of greater than a threshold are selected (highlighted in black).

We then applied PCA on the subset of 3,792 highly variable genes and used JackStraw^7, 18^ to identify significant PCs. We identified 32 significant PCs. To identify PCs representing non-RGC markers, we found 68 genes (e.g., Gpr37, Gldc, and Epas1) from literature that are known markers for major non-RGC retinal cell types including Muller, Fibroblast, Astrocyte, Amacrine, Endothelium, Horizontal, Photoreceptor, and Microglia cells. We then computed the pairwise correlation between any of these 68 markers and any of 32 PCs. We then excluded those PCs that showed a significant correlation (Pearson) with known non-RGC markers. The first PC had the highest correlation of 0.58 with several non-RGC markers and was excluded from the downstream analysis.

Graph-based clustering^25,26^ with SLM modularity optimization^27^ identified 24 clusters visualized using tSNE.^19^Figure 16 shows the tSNE plot of the identified clusters of cells color coded and numbered based on the labels identified by SLM algorithm.

**Figure 16.**
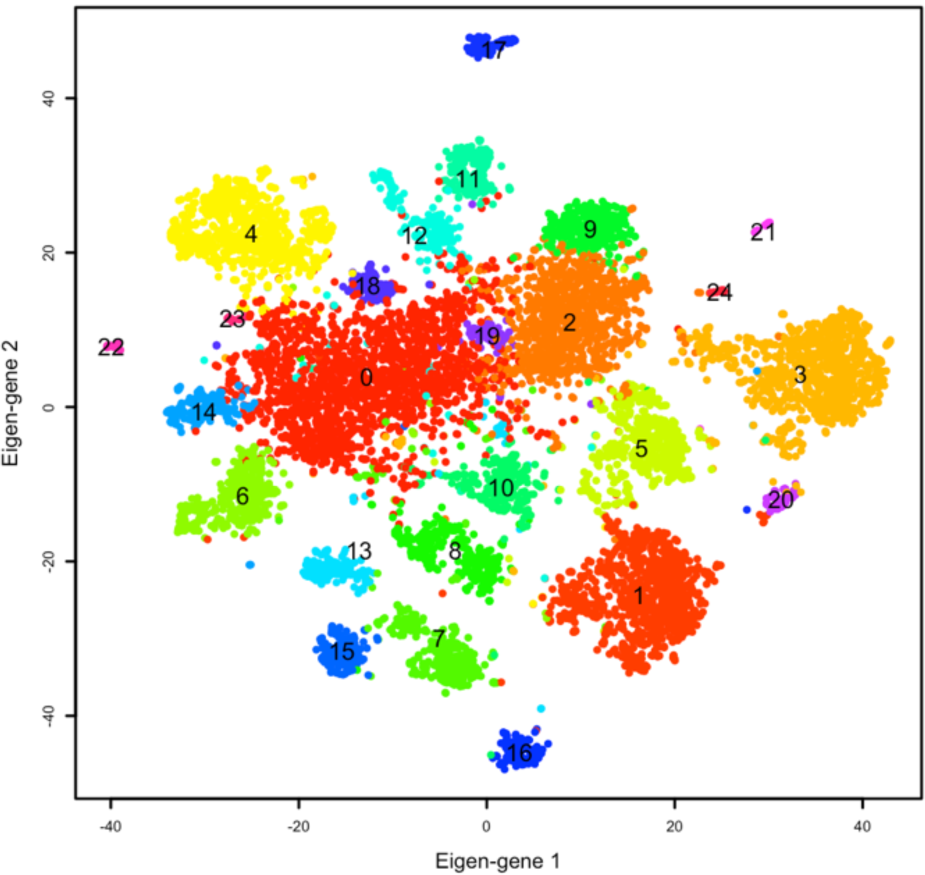
Final clustering identified 24 clusters of cells. Contaminating clusters of non-RGC cells will be identified and excluded.

To identify potential non-RGC cells in the tSNE space, we first identified marker genes differentially expressed in each cluster by comparing genes expressed in each cluster with genes expressed in the remaining clusters using likelihood-ratio.^21,28^ Post hoc analysis to qualitatively and quantitively identifying biological attribute of clusters, particularly those with significant expression of non-RGC markers, identified a few closers of non-RGC cells.

Figure 17 shows the heatmap of genes that were differentially expressed in 24 clusters in a single snapshot and figure 18 presents sample genes that were highly selective in terms of expression in different clusters.

**Figure 17.**
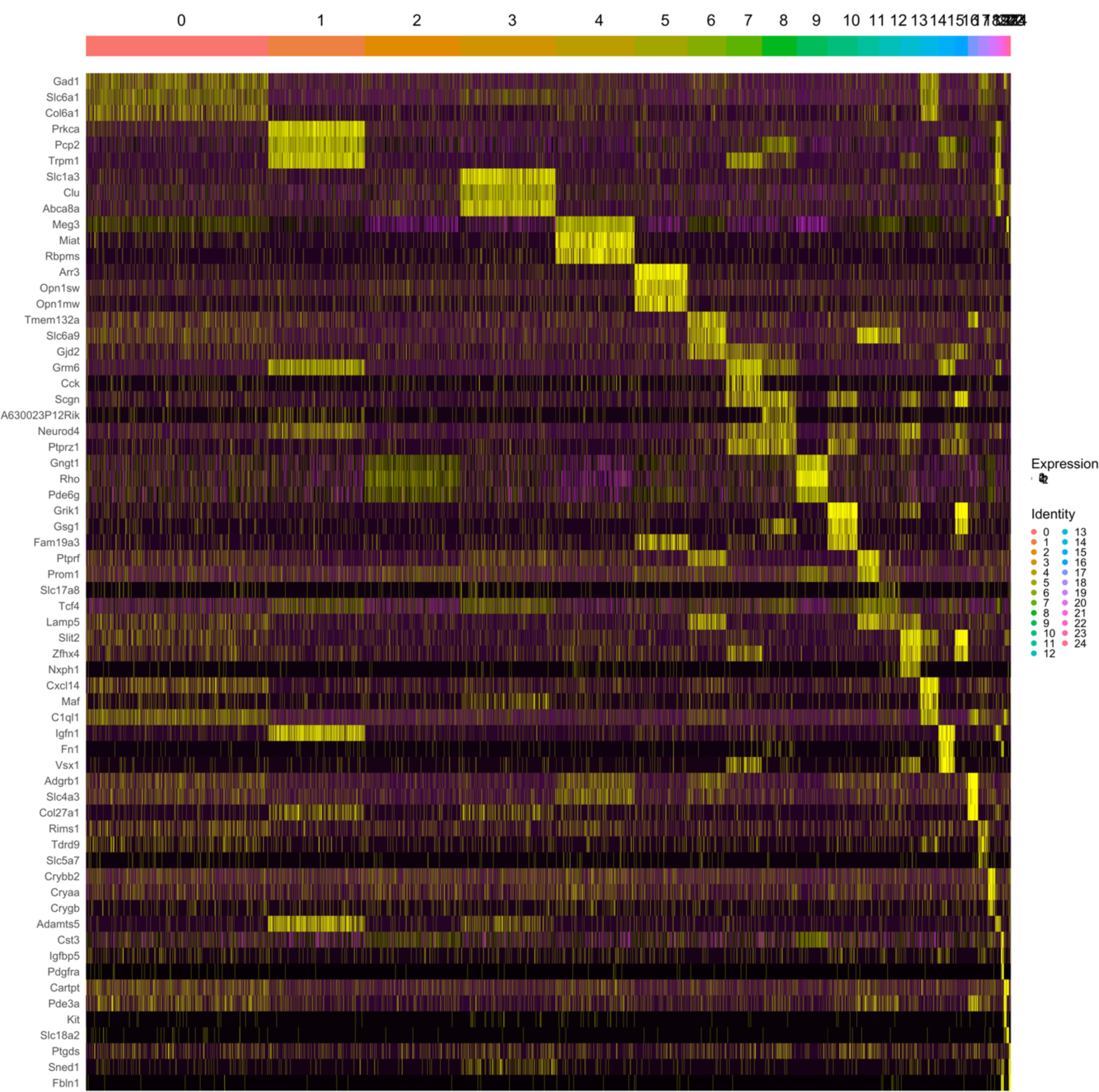
Heatmap plot of genes expressed in different clusters. Cells are sorted based on cluster identity as shown on top of the plot. Genes that were differentially expressed are presented on the y axis. Yellow color shows high expression.

**Figure 18.**
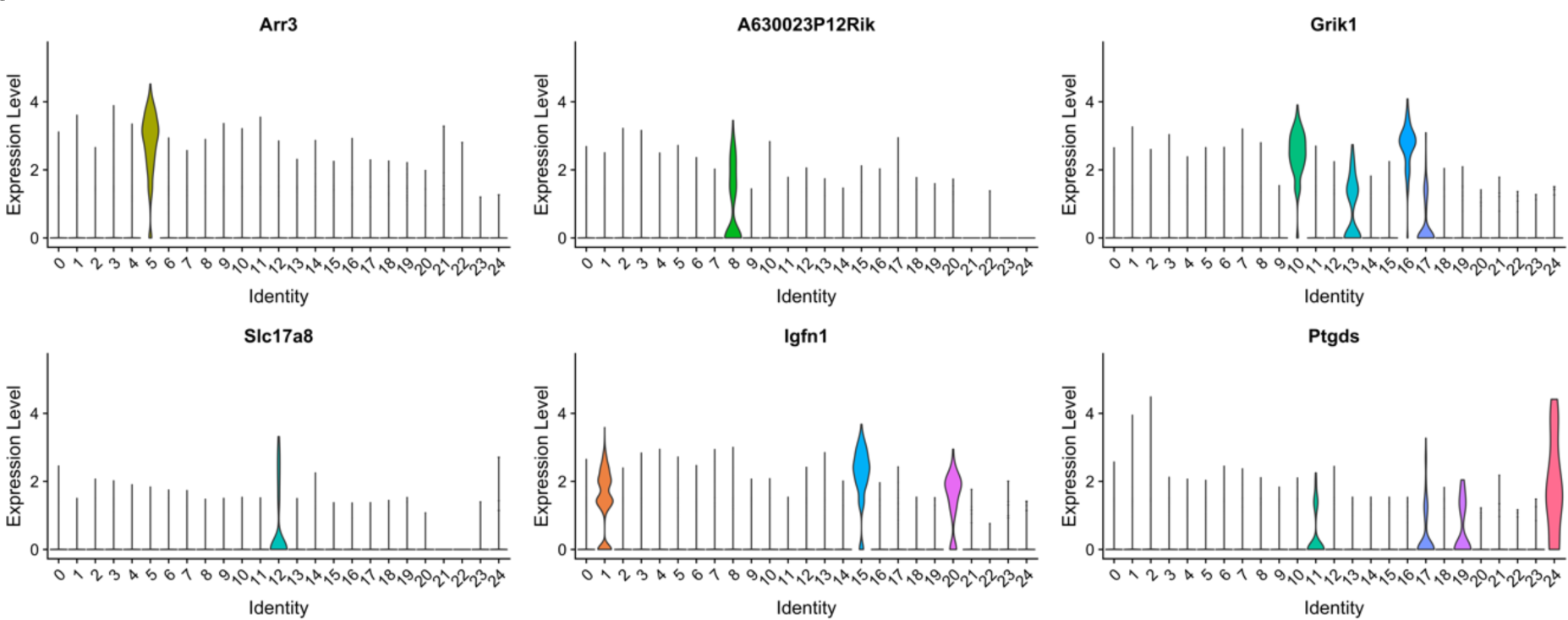
Differentially expressed genes. Six sample differentially expressed marker genes for several clusters computed suing likelihood-ration.

After posthoc analysis, we identified that 5 out of 14 genes differentially expressed in cluster number 5 were known photoreceptor markers. Therefore, cells in this cluster were excluded from the analysis. This process, coupled with domain knowledge, can be performed iteratively to exclude other potential non-RGC cells from the analysis.

### Single ventral midbrain cells

Eight DBA/2J or DBA/2J-Gpnmb mice in an age range of 7-8 days old were anesthetized and ventral midbrain tissues were removed and dissociated gently. Fluidigm C1-HT microfluidics plates were used to isolate and generate scRNA-seq libraries of full-length mRNAs using SMART-Seq v4. Three libraries each one including 400 cells were sequenced using HiSeq3000, PE151. Following alignment using STAR, expression was normalized to log2(FPKM+1) across ∼25,000 unique transcript models. Each cell was sequenced to a depth of ∼490,000 with an average of 1,778 genes per cell.

To enhance the power of unsupervised clustering for discovering RGC subtypes, we filtered cells expressing fewer than 200 genes and genes expressed in fewer than five cells. A total of 1,149 cells expressing 14,147 genes were included for the downstream analysis. Figure 19 shows the histogram of the number of genes expressed in cells. In average, 1,778 genes were expressed in cells.

**Figure 19.**
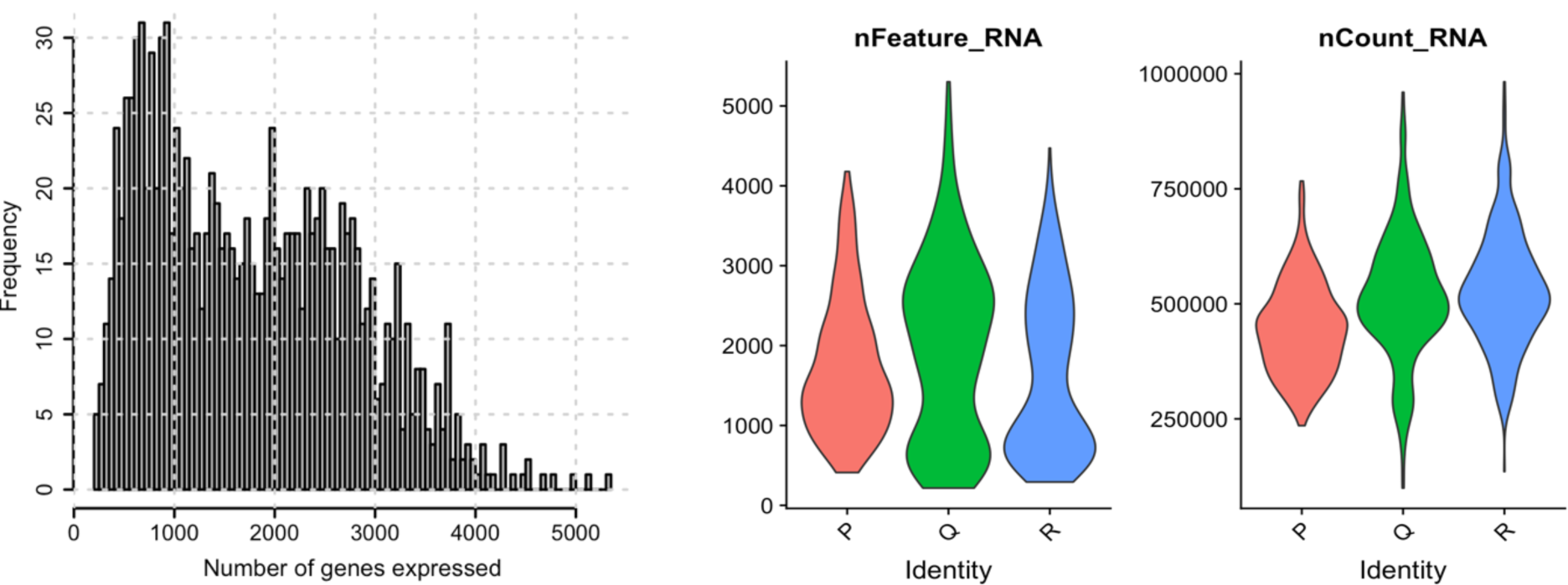
Number of genes expressed in ventral midbrain cells. Left panel shows the histogram of number of genes expressed in all cells. Middle panel shows the violin plot of number of genes expressed in three plates, each plate with 400 ventral midbrain cells and the right panel represents the number fragments per kilobase million (FPKM) in three plates each with 400 ventral midbrain cells.

We then scaled and centered the data along each gene then selected highly variable genes by computing average expression and dispersion of each gene. We identified 5,516 highly variable genes based on average expression and dispersion (Fig. 20). Reducing the number of genes reduces the computational complexity and improves the ability to identify selective genes of different midbrain cell types.

**Figure 20.**
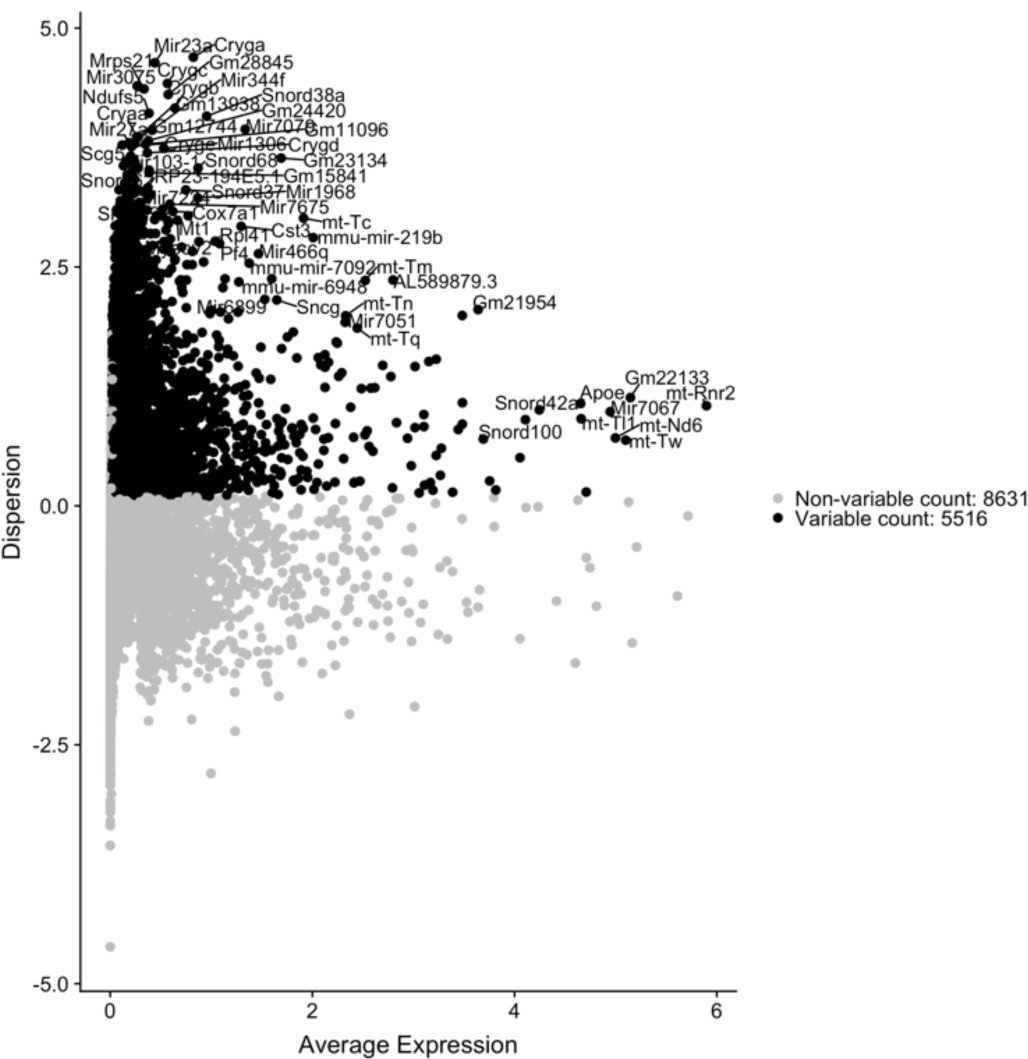
Average expression versus dispersion of genes in our dataset. Genes that having an average expression and dispersion of greater than a threshold are selected (represented in black). Names of some of the highly variable genes are provided.

We then applied PCA on the subset of 5,516 highly variable genes in order to capturing the primary structures and patterns in the transcriptome data. This process generates 5,516 principal components, however, only a small number of these components capture the variance exist in the data. We used JackStraw^7, 18^ method and identified 34 significant PCs.

We then performed graph-based clustering that starts with a K-nearest neighbor (KNN) graph, with edges drawn between cells with similar gene expression patterns, and then partitioned this graph into highly interconnected quasi-cliques or communities, as outlined in previous publications.^25 26^ We then applied modularity optimization techniques proposed in Louvain algorithm or SLM ^27^, to iteratively group cells together, with the goal of optimizing the standard modularity function. We then visualized the clusters using tSNE. ^19^ We set the effective number of neighbors in tSNE represented by “perplexity” parameter to 30. In fact, cells with similar gene expression patterns will fall closely, and hence distinct cell types should form clusters in the 2-dimensional tSNE space. Figure 6 shows the tSNE plots of the cells we identified. We identified 13 different clusters of cells (Fig. 20).

**Figure 21.**
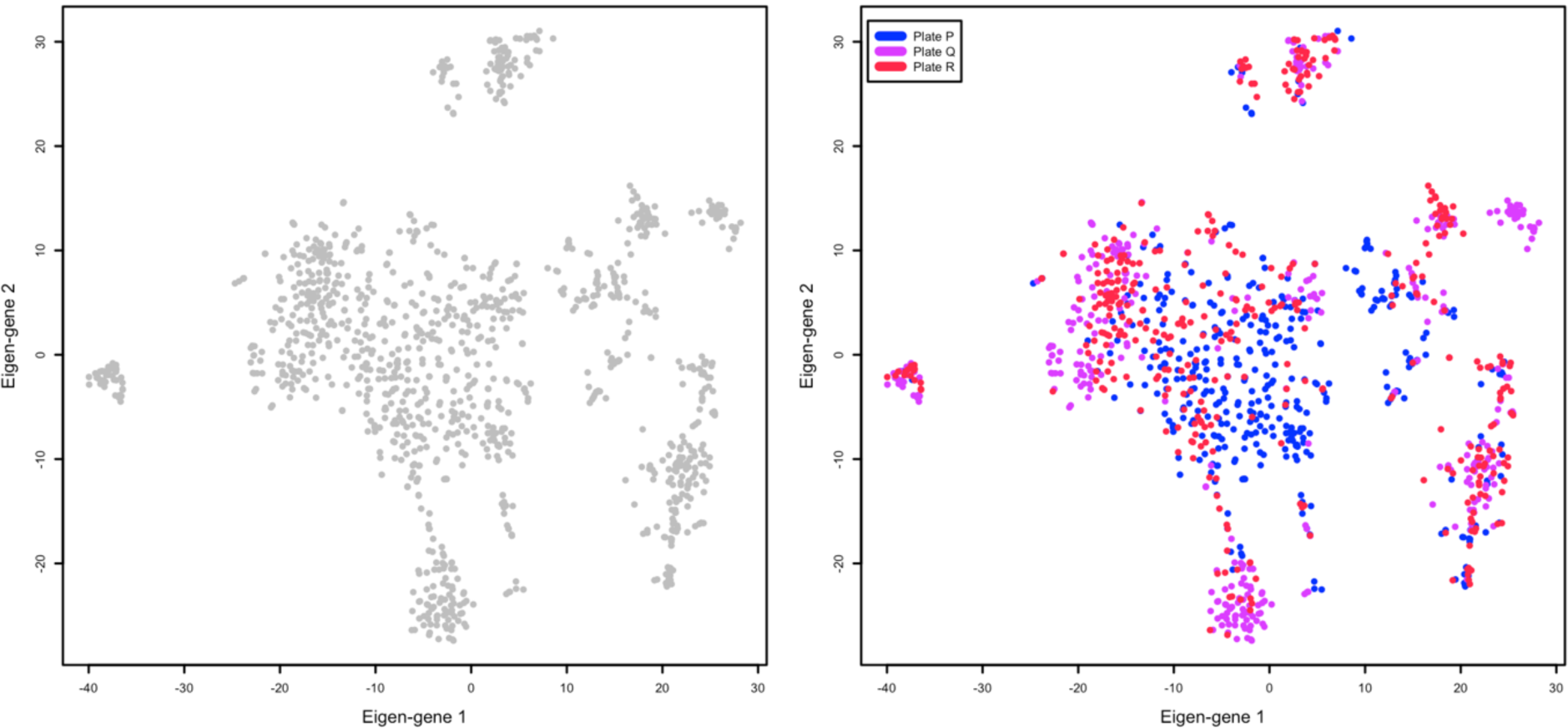
Cell mapped on 2-D tSNE space. Cells with similar transcriptome patterns have been grouped together. Left panel shows all cells and right panel shows cells color coded by the corresponding plate (batch) in the experimental workflow.

We identified marker genes differentially expressed in each cluster by comparing genes expressed in each cluster with genes expressed in the remaining clusters. More specifically, we required a gene in each cluster to be detected at a minimum percentage and a minimum expression level compared to genes in the rest of clusters. This was investigated using likelihood-ratio test for single cell gene expression.^21, 28^ We explored other techniques including negative binomial generalized linear model, and negative binomial distribution implemented in the DESeq2 algorithm^29^ to assure repeatability.

**Figure 22.**
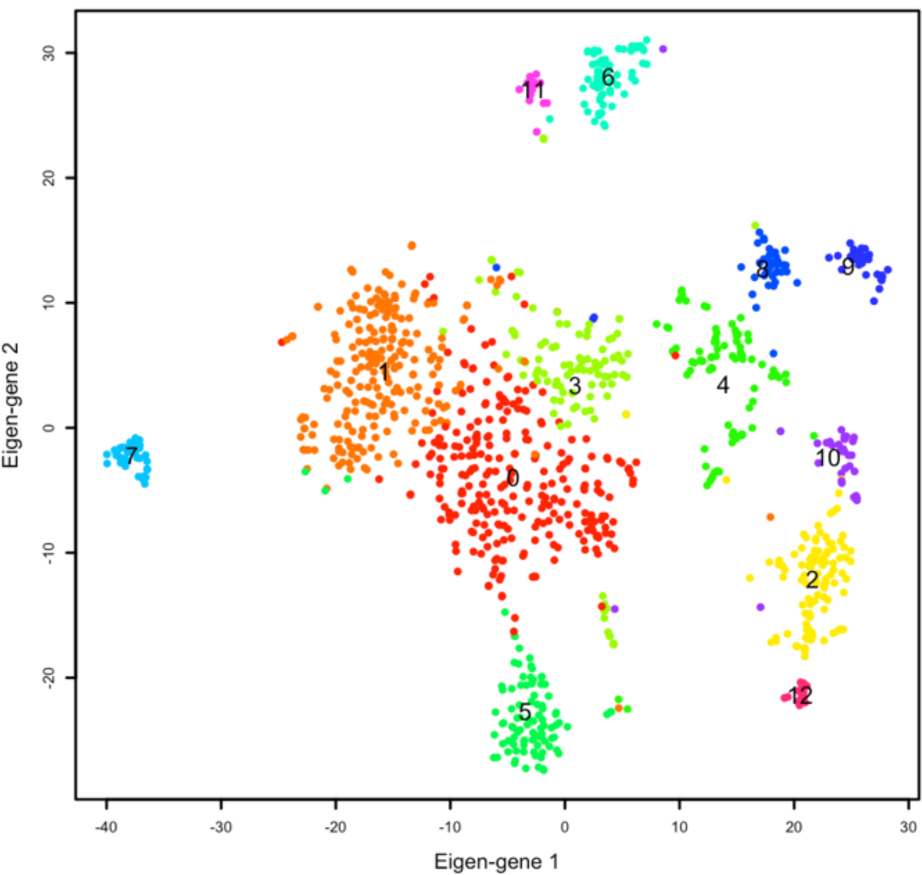
Final clusters. We identified 13 different clusters of ventral midbrain cells.

We subsequently performed supplementary *post hoc* qualitative analysis to identify biological attribute of clusters. Specifically, we identified clusters of cells that co-express known ventral midbrain markers indicated before. This is a classical approach and has been used in the latest studies aiming at identifying different cell types. We identified ten genes differentially expressed in our dataset that were matching with La Manno el al.^31^ : Pdgfra, Clu, Ccl4, Cst3, Gad2, Six3, Nefl, Rit2, Cd36, and Apod (Figure 23).

**Figure 23.**
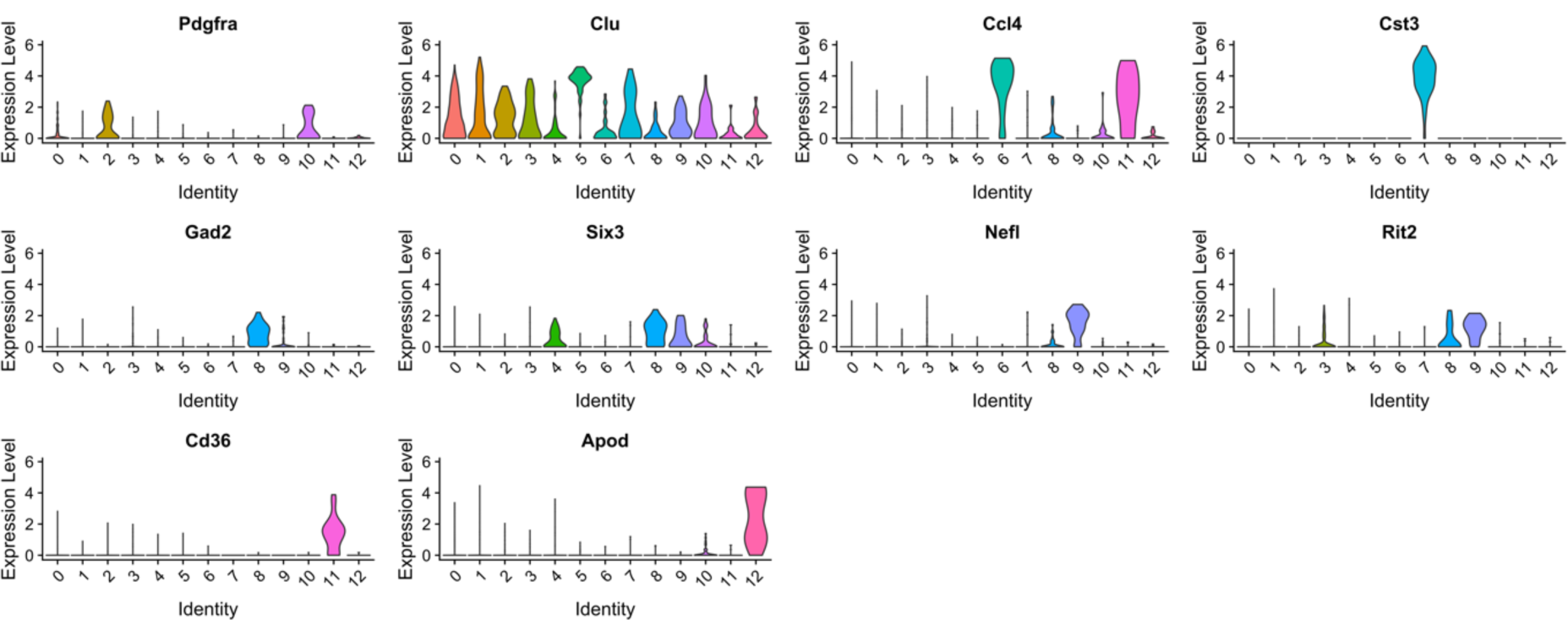
Differentially expressed genes. Ten sample differentially expressed genes for ventral midbrain cells identified using likelihood-ratio.

Figure 24 shows a snapshot plot of all cells, sorted by cluster membership, versus top genes expressed in clusters. Yellowish color shows high expression of those genes in different clusters.

**Figure 24.**
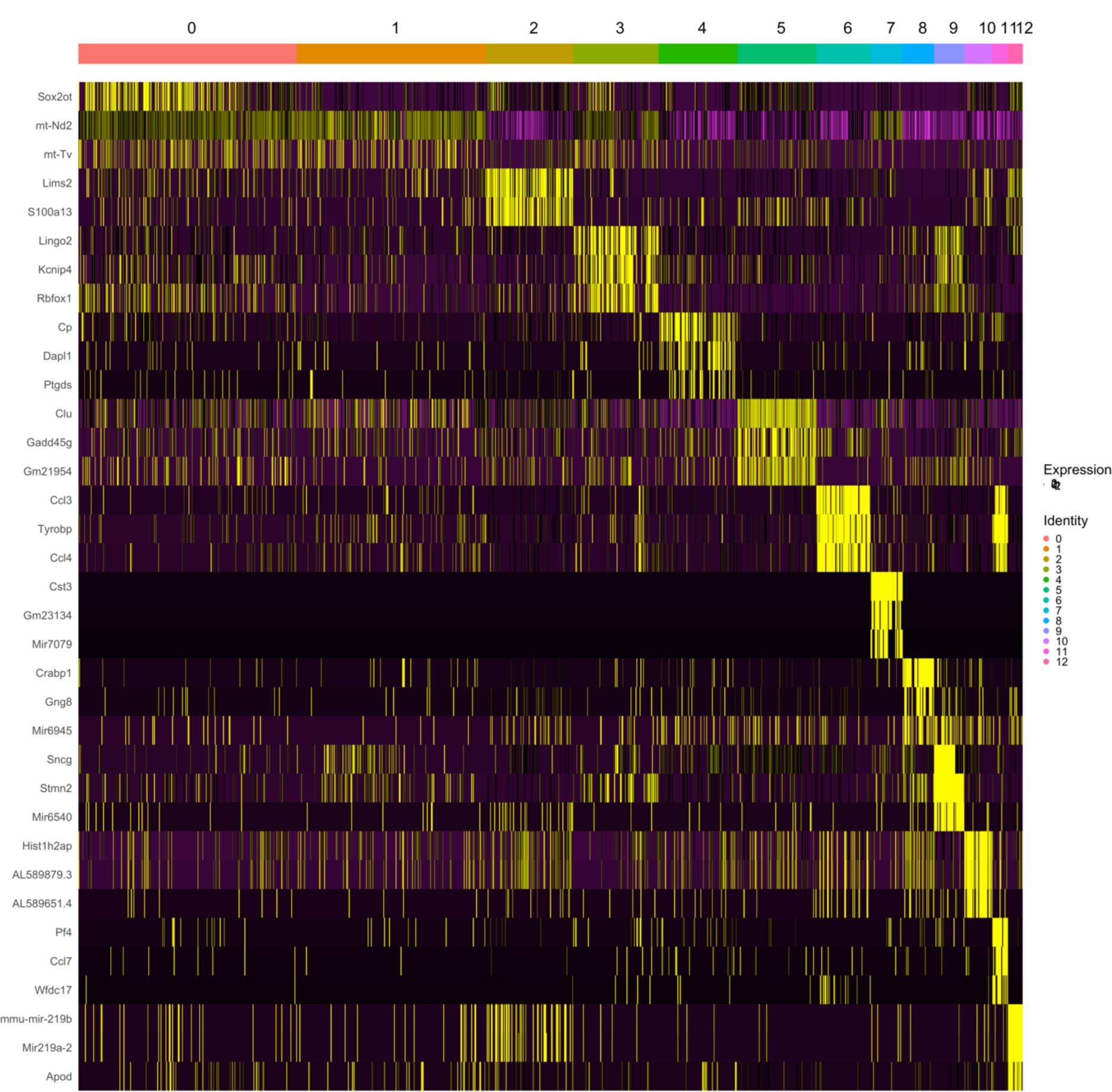
Heatmap plot of top genes expressed in different clusters. Cells are sorted based on cluster identity as shown on top of the plot. Genes that were differentially expressed are presented on the y axis. Yellow color shows high expression.

## Acknowledgement

This study was supported by a Stein Innovation Award from Research to Prevent Blindness and Center for Integrative and Translational Genomics (CITG) and in part by the Bright Focus Foundation.

## Notes

**Grant Support**: Supported in part by grants of Stein Innovation Award from Research to Prevent Blindness and Center for Integrative and Translational Genomics (CITG).

### Competing Interest Statement

The authors have declared no competing interest.

### Summary of Updates

Complete names of authors, affiliations, and acknowledgement.

## References

1. Huberman AD, Wei W, Elstrott J, Stafford BK, Feller MB, Barres BA. Genetic identification of an On-Off direction-selective retinal ganglion cell subtype reveals a layer-specific subcortical map of posterior motion. Neuron 2009;62:327–334.

2. Rousso DL, Qiao M, Kagan RD, Yamagata M, Palmiter RD, Sanes JR. Two Pairs of ON and OFF Retinal Ganglion Cells Are Defined by Intersectional Patterns of Transcription Factor Expression. Cell Rep 2016;15:1930–1944.

3. Sweeney NT, James KN, Nistorica A, Lorig-Roach RM, Feldheim DA. Expression of transcription factors divides retinal ganglion cells into distinct classes. J Comp Neurol 2017.

4. Regev A, Teichmann SA, Lander ES, et al. The Human Cell Atlas. Elife 2017;6.

5. Levin LA. Retinal ganglion cells and supporting elements in culture. J Glaucoma 2005;14:305–307.

6. Rokicki W, Dorecka M, Romaniuk W. [Retinal ganglion cells death in glaucoma--mechanism and potential treatment. Part II]. Klin Oczna 2007;109:353–355.

7. Masland RH. The fundamental plan of the retina. Nat Neurosci 2001;4:877–886.

8. Demb JB. Cellular mechanisms for direction selectivity in the retina. Neuron 2007;55:179–186.

9. Struebing FL, Lee RK, Williams RW, Geisert EE. Genetic Networks in Mouse Retinal Ganglion Cells. Front Genet 2016;7:169.

10. Sumbul U, Song S, McCulloch K, et al. A genetic and computational approach to structurally classify neuronal types. Nat Commun 2014;5:3512.

11. Volgyi B, Chheda S, Bloomfield SA. Tracer coupling patterns of the ganglion cell subtypes in the mouse retina. J Comp Neurol 2009;512:664–687.

12. Coombs J, van der List D, Wang GY, Chalupa LM. Morphological properties of mouse retinal ganglion cells. Neuroscience 2006;140:123–136.

13. Kong JH, Fish DR, Rockhill RL, Masland RH. Diversity of ganglion cells in the mouse retina: unsupervised morphological classification and its limits. J Comp Neurol 2005;489:293–310.

14. Tanabe Y, Jessell TM. Diversity and pattern in the developing spinal cord. Science 1996;274:1115–1123.

15. Satija R, Farrell JA, Gennert D, Schier AF, Regev A. Spatial reconstruction of single-cell gene expression data. Nat Biotechnol 2015;33:495–502.

16. Butler A, Hoffman P, Smibert P, Papalexi E, Satija R. Integrating single-cell transcriptomic data across different conditions, technologies, and species. Nat Biotechnol 2018;36:411–420.

17. Leek JT, Storey JD. Capturing heterogeneity in gene expression studies by surrogate variable analysis. PLoS Genet 2007;3:1724–1735.

18. Chung NC, Storey JD. Statistical significance of variables driving systematic variation in high-dimensional data. Bioinformatics 2015;31:545–554.

19. Laurens van der Maaten GH. Visualizing Data using t-SNE. Journal of Machine Learning Research 2008;9:2579 – 2605.

20. Martin Ester H-PK, Jörg Sander, Xiaowei Xu. A density-based algorithm for discovering clusters in large spatial databases with noise Knowledge discovery and data mining (KDD) 1996;226--231.

21. McDavid A, Finak G, Chattopadyay PK, et al. Data exploration, quality control and testing in single-cell qPCR-based gene expression experiments. Bioinformatics 2013;29:461–467.

22. Macosko EZ, Basu A, Satija R, et al. Highly Parallel Genome-wide Expression Profiling of Individual Cells Using Nanoliter Droplets. Cell 2015;161:1202–1214.

23. Pandey S, Shekhar K, Regev A, Schier AF. Comprehensive Identification and Spatial Mapping of Habenular Neuronal Types Using Single-Cell RNA-Seq. Curr Biol 2018;28:1052–1065 e1057.

24. Shekhar K, Lapan SW, Whitney IE, et al. Comprehensive Classification of Retinal Bipolar Neurons by Single-Cell Transcriptomics. Cell 2016;166:1308–1323 e1330.

25. Xu C, Su Z. Identification of cell types from single-cell transcriptomes using a novel clustering method. Bioinformatics 2015;31:1974–1980.

26. Levine JH, Simonds EF, Bendall SC, et al. Data-Driven Phenotypic Dissection of AML Reveals Progenitor-like Cells that Correlate with Prognosis. Cell 2015;162:184–197.

27. Blondel VD, Guillaume J-L, Lambiotte R, Lefebvre E. Fast unfolding of communities in large networks. Journal of Statistical Mechanics: Theory and Experiment 2008;2008:P10008.

28. McDavid A, Dennis L, Danaher P, et al. Modeling bi-modality improves characterization of cell cycle on gene expression in single cells. PLoS Comput Biol 2014;10:e1003696.

29. Love MI, Huber W, Anders S. Moderated estimation of fold change and dispersion for RNA-seq data with DESeq2. Genome Biol 2014;15:550.

30. Zheng GX, Terry JM, Belgrader P, et al. Massively parallel digital transcriptional profiling of single cells. Nat Commun 2017;8:14049.

31. La Manno G, Gyllborg D, Codeluppi S, et al. Molecular Diversity of Midbrain Development in Mouse, Human, and Stem Cells. Cell. 2016;167(2):566–580 e519.

